# Long-range functional coupling predicts performance: oscillatory EEG networks in multisensory processing

**DOI:** 10.1101/014423

**Authors:** Peng Wang, Florian Göschl, Uwe Friese, Peter König, Andreas K. Engel

**Affiliations:** Department of Neurophysiology and Pathophysiology, University Medical Center Hamburg-Eppendorf, Martinistr. 52, 20246 Hamburg, Germany; Institute of Cognitive Science, University of Osnabrück, Albrechtstr. 28, 49069 Osnabrück, Germany

**Author notes:** Correspondence address: Peng Wang, Florian Göschl, Department of Neurophysiology and Pathophysiology, University Medical Center Hamburg-Eppendorf, Martinistr. 52, 20246 Hamburg, Germany. These authors contributed equally (Peng Wang, Florian Göschl).

**Keywords:** oscillations, multisensory, EEG, phase locking value, phase coupling, phase-amplitude coupling, attention

## Abstract

The integration of sensory signals from different modalities requires flexible interaction of remote brain areas. One candidate mechanism to establish communication in the brain is transient synchronization of oscillatory neural signals. Although there is abundant evidence for the involvement of cortical oscillations in brain functions based on the analysis of local power, assessment of the phase dynamics among spatially distributed neuronal populations and their relevance for behavior is still sparse. In the present study, we investigated the interaction between remote brain areas by analyzing high-density electroencephalogram (EEG) data obtained from human participants engaged in a visuotactile pattern matching task. We deployed an approach for purely data-driven clustering of neuronal phase coupling in source space, which allowed imaging of large-scale functional networks in space, time and frequency without defining a priori constraints. Based on the phase coupling results, we further explored how brain areas interacted across frequencies by computing phase-amplitude coupling. Several networks of interacting sources were identified with our approach, synchronizing their activity within and across the theta (~5 Hz), alpha (~10 Hz), and beta (~ 20 Hz) frequency bands and involving multiple brain areas that have previously been associated with attention and motor control. We demonstrate the functional relevance of these networks by showing that phase delays – in contrast to spectral power – were predictive of task performance. The data-driven analysis approach employed in the current study allowed an unbiased examination of functional brain networks based on EEG source level connectivity data. Showcased for multisensory processing, our results provide evidence that large-scale neuronal coupling is vital to long-range communication in the human brain and relevant for the behavioral outcome in a cognitive task.

## Introduction

The world surrounding us is inherently multimodal and requires continuous processing and accurate combination of information from the different sensory systems. Constantly, agreement or conflict of different sensory signals is evaluated in the human brain, requiring flexible interactions of functionally specialized and spatially distributed areas. Crossmodal interplay has been shown to impact perception (McGurk and MacDonald, 1976) as well as a broad range of cognitive processes (Driver and Spence, 2000; Macaluso and Driver, 2005; Stein, 2012). Yet, the neurophysiological implementation of long-range interactions in the brain remains a subject of active exploration. It has been proposed that synchronization of oscillatory signals might underlie the dynamic formation of task-dependent functional networks (Engel et al., 2001; Fries, 2005; Salinas and Sejnowski, 2001; Varela et al., 2001; Womelsdorf et al., 2007). More recently, transient synchronization of neuronal signals has also been implicated in establishing relationships across different sensory systems (Kayser et al., 2008; Lakatos et al., 2007), allowing the preferential routing of matching crossmodal information to downstream assemblies (Senkowski et al., 2008). In spite of ample evidence supporting the importance of cortical oscillations as measured by local spectral power in multisensory processing (Bauer et al., 2009; Gleiss and Kayser, 2014; Göschl et al., 2015; Schneider et al., 2008; Schneider et al., 2011; Senkowski et al., 2014; for reviews see Keil and Senkowski, 2018 and Senkowski et al., 2008), studies investigating large-scale functional connectivity based on whole-brain measurements such as EEG or magnetoencephalography (MEG) are still rare. Importantly, local signal power may convey different information as measures of neural interaction (Schneidman et al., 2006; Snyder et al., 2015). What is more, first order measures (like spectral power of single nodes) are not adequate for a comprehensive description of network properties, which asks for at least second order (i.e. pairwise) measures (Schneidman et al., 2006). Thus, it seems appropriate to make use of measures that allow for illustrating global patterns of neural communication involving local as well as long-range connections. One candidate for such a holistic measure is phase coupling of multiple brain signals, as assessed for example by phase locking values (PLVs). Experimental evidence directly linking neural coupling and multimodal processing comes from invasive recordings in behaving animals (Kayser and Logothetis, 2009; Maier et al., 2008; von Stein et al., 2000) as well as a small number of studies in humans performed with EEG and MEG (Doesburg et al., 2008; Hummel and Gerloff, 2005; Lange et al., 2013; van Driel et al., 2014). The studies available so far have relied on specific a priori assumptions regarding brain locations, time windows, or oscillation frequencies of the involved networks and consequently only reflect a fraction of the actual network dynamics.

In the present study, we sought to investigate global patterns of neuronal coupling in multisensory processing without predefined constraints and to characterize the involved networks in space, time and frequency. To do so, we adapted an approach for purely data-driven analysis of neuronal coupling in source space that has recently been developed by our group (Friese et al., 2016; Hipp et al., 2011). We sought to assess functional connectivity during the processing of crossmodal sensory input, using measures of phase coupling as well as phase-amplitude coupling. Connectivity indices were submitted to a multi-dimensional clustering approach with the full range of time, frequency and cortical space, to detect functional networks involved in our cognitive task. Specifically, we investigated whether coupling of oscillatory signals across cortical regions was crucial for the perception and integration of multimodal information by directly linking the coupling characteristics of the identified networks to participants’ task performance.

## Materials and Methods

The dataset reported on in this manuscript was published earlier (Göschl et al., 2015), using a different analysis approach and research purpose.

### Participants

16 right-handed volunteers (12 females, mean age 25.4, range 21–33) participated in the current experiment and were compensated in monetary form. Participants had normal or corrected to normal vision and reported no history of neurological or psychiatric illness. The study was approved by the Ethics Committee of the Medical Association Hamburg and conducted in accordance with the Declaration of Helsinki. Prior to the recordings, all participants provided written informed consent.

### Task design

In the current experiment, we employed a setup similar to the one used in a previous behavioral study (Göschl et al., 2014). Figure 1 provides an overview of events and timing of the visuotactile matching paradigm used here. The stimulus set consisted of four spatial patterns, each formed by three dots (Figure 1A). Stimulation was always bimodal, with visual and tactile patterns being presented concurrently on a computer screen and to participants’ index fingertip via a Braille stimulator (QuaeroSys Medical Devices, Schotten, Germany). Visual stimuli appeared left of a central fixation cross and were embedded in a noisy background while tactile patterns were delivered to the right index finger. Stimulus duration was 300 ms for both patterns. To familiarize participants with the tactile stimuli, we conducted a delayed-match-to-sample training task prior to the actual experiment (for details see Göschl et al., 2014). One participant was excluded after the training procedure due to insufficient performance.

**Figure 1.**
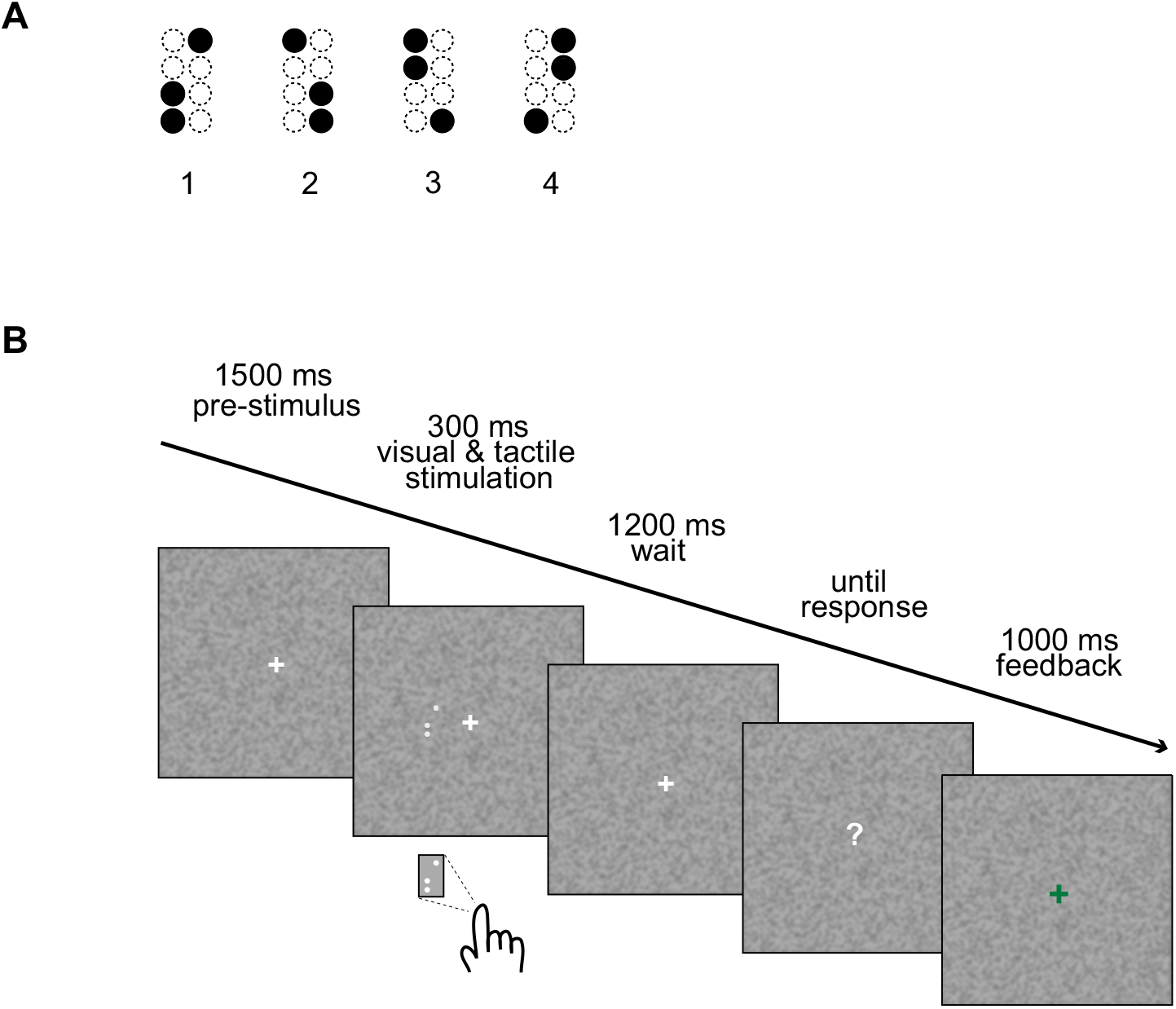
Schematic representation of the visuotactile detection task. (A) The four stimulus patterns used in our experiment. (B) The trial sequence. After a pre-stimulus interval of 1500 ms, visual and tactile stimuli were presented simultaneously for 300 ms, followed by a wait interval of 1200 ms. After that, a question mark appeared on the screen indicating that responses could be given. After a button press, visual feedback was given (1000 ms; adopted from Göschl et al., 2014, 2015).

Participants were instructed to detect predefined target stimuli that could appear in both modalities. At the beginning of each experimental block, one of the four patterns was defined as the target stimulus (the other three patterns were non-targets, respectively) by simultaneously presenting it on the computer screen and by means of the Braille stimulator (four times). During the experimental block, the target could appear in the visual or the tactile modality alone, in both or in neither of the two. Participants were asked to decide whether the presented stimuli matched the previously defined target or not and press one of two response buttons accordingly. Participants responded with their left hand via button press on a response box (Cedrus, RB-420 Model, San Pedro, USA) and visual feedback (a green ‘+’ or a red ‘–’) was given in every trial. The timing of events is displayed in Figure 1B.

In two sessions taking place within three days, we recorded 1152 trials from each participant. The design was counterbalanced with respect to crossmodal stimulus congruence, target definition and presentation frequency of each of the four patterns (for details, see Göschl et al., 2014). Data from the two recording sessions were pooled and trials grouped according to target appearance, resulting in the following conditions: tactile-matching only (a tactile target presented with a visual non-target; labeled as T, 16.7%), visual-matching only (a visual target appearing with a tactile non-target; V, 16.7%), and visual-tactile targets (VT, 16.7%) as well as non-target congruent (C, 16.7%) or incongruent pairs (IC, 33.3%). Unless specified otherwise, we focus our analysis on data from correctly answered trials. The number of trials was stratified across conditions (i.e. the same number of trials per condition entered the analysis).

Key mapping (for ‘target’ and ‘non-target’-buttons) was counterbalanced across participants and sessions. Sounds associated with pin movement in the Braille cells were masked with pink noise administered via foam-protected air tube earphones at 75 dB sound pressure level (Eartone, EAR Auditory Systems, AearoCompany). We used Presentation software (Neurobehavioral Systems, version 16.3) to control stimulus presentation and to record participants’ response times (RT) and accuracies.

### EEG data collection and preprocessing

Electroencephalographic data were acquired from 126 scalp sites using Ag/AgCl ring electrodes mounted into an elastic cap (EASYCAP, Herrsching, Germany). Two additional electrodes were placed below the eyes to record the electrooculogram. EEG data were recorded with a passband of 0.016-250 Hz and digitized with a sampling rate of 1000 Hz using BrainAmp amplifiers (BrainProducts, Munich, Germany). During the recordings, the tip of the nose served as a reference but subsequently we re-referenced the data to common average. Preprocessing of the EEG data was carried out in Matlab 8.0 (MathWorks, Natick, MA) using custom-made scripts, as well as routines incorporated in EEGLAB 11.0 (Delorme and Makeig, 2004; http://sccn.ucsd.edu/eeglab/). Offline, the data were band-pass filtered (0.3-180 Hz), down-sampled to 500 Hz and epoched from – 400 to + 1400 ms around stimulus onset. Next, all trials were inspected visually and those containing muscle artifacts were rejected. To remove artifacts related to eyeblinks, horizontal eye movements and electrocardiographic activity, we applied an independent component analysis (ICA) approach (Hipp and Siegel, 2013). Furthermore, we employed the COSTRAP algorithm (correction of saccade-related transient potentials; Hassler et al., 2011) to control for miniature saccadic artifacts. This algorithm has been used in previous studies (e.g. Friese et al., 2013; Hassler et al., 2013) to suppress ocular sources of high frequency signals. Employing this multilevel artifact correction procedure, 88% of all recorded trials (range: 75% to 95% for individual participants) were retained. The number of trials for all conditions was stratified before applying the network identification approach based on differences between conditions.

### Source estimation of frequency-specific activity

We constructed our source space based on the template brain ICBM152 from the Montreal Neurological Institute (Mazziotta et al., 1995). The cortex of this brain template was segmented with freesurfer software (Dale et al., 1999; Fischl et al., 1999). To generate the source space, we started from an icosahedron and then recursively subdivided its edges, the middle point of which would be repositioned to its circumscribed sphere as new vertices. Then the circumscribed sphere, with 162 vertices on it, was morphed onto the outer surface of the white matter (i.e. the inner surface of the gray matter) of each hemisphere, which yielded 324 locations homogeneously covering the whole cortex. By excluding locations which were unlikely to elicit significant brain responses (e.g., the corpus callosum), we reduced the number of source locations to 300 (Figure S1). This procedure was accomplished with the MNE software package (Gramfort et al., 2014). We chose a relatively low number of source locations for reasons of computational efficiency, and in order to be compatible with the physical spatial resolution of the EEG sensors. In a next step, we computed leadfields with these source locations and averaged sensor positions from 30 subjects, whose data were registered in previous studies in our laboratory, using the very same EEG channel layout. Leadfield calculation was realized with a 3-shell head model (Nolte and Dassios, 2005). We did not apply compensation for ICA component reduction, with the assumption that the leadfield distribution was not altered by removing components related to eye movements or other muscle activity (Hipp and Siegel, 2015). Making use of linearly constrained minimum variance (LCMV) methods (Van Veen et al., 1997), we projected the original sensor-level waveforms to the source locations defined above. For each source location, we computed three orthogonal filters (one for each spatial dimension) that passed activity from the location of interest with unit gain while maximally suppressing activity from all other sources. Next, we linearly combined the three filters into a single filter in the direction of maximal variance (Hipp et al., 2011). In order to avoid spurious effects resulting from unequal filters in between-condition comparisons, we used data from all conditions (with the same number of trials for each condition) to compute a common filter. To derive source estimates, we multiplied the cleaned sensor data with the real-valued filter. High source correlations can reduce source amplitudes estimated with beamforming due to source cancelation (Hipp et al., 2011; Van Veen et al., 1997) which might influence the magnitude of cortico-cortical coupling estimates. However, within the range of physiological source correlations (Leopold et al., 2003), the identification of cortico-cortical coupling using the beamforming method should be possible (Kujala et al., 2008). Moreover, although source cancelation might reduce the sensitivity to detect coupling, it should not lead to false positive results.

### EEG time-frequency decomposition

The time-frequency decomposition was performed in cortical source space. Since we were interested in induced activities rather than amplitude changes time-locked to the stimulus, we removed the event related potentials (ERP) from each data trial prior to the spectral analysis. All spectral estimates were performed with the multitaper method (Mitra and Pesaran, 1999; Percival and Andrew, 1993) and computed across 21 logarithmically scaled frequencies from 4 to 128 Hz with 0.25 octave steps and across 17 time points from – 300 to 1300 ms in steps of 100 ms. The temporal and spectral smoothing was performed as follows. For frequencies larger or equal to 16 Hz, we used temporal windows of 250 ms and adjusted the number of slepian tapers to approximate a spectral smoothing of 3/4 octave; for frequencies lower than 16 Hz, we adjusted the time window to yield a frequency smoothing of 3/4 octaves with a single taper (Hipp et al., 2011). In case the window extended outside of the trial range (– 400 to 1400 ms), zeros were padded. We characterized power and phase locking responses relative to the prestimulus baseline using the bin centered at – 200 ms. The employed time-frequency transformation ensured a homogenous sampling and smoothing in time and frequency, as required for subsequent network identification within this multi-dimensional space.

### Phase-phase coupling analysis

To quantify the frequency-dependent synchronization in source space, we estimated phase locking values (PLV; (Lachaux et al., 1999)) – a measure less dependent on changes in signal amplitude than coherence. Anyway, using coherence measures yielded similar results (data not shown).

The PLV between pairs of signals *X*(*f, t*) and *Y*(*f, t*) was computed according to the following equation:

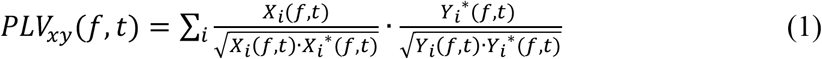

Here *i* is the trial index, *t* is the time bin index, and *f* is the frequency bin index.

Since PLV is positively biased to 1 with decreasing number of independent spectral estimates (degrees of freedom), we stratified the sample size and used the same number of trials for the comparison between conditions. We also performed a nonlinear transform (Jarvis and Mitra, 2001) with the following equation to render the PLV distribution to approximately Gaussian before submitting the values to network identification:

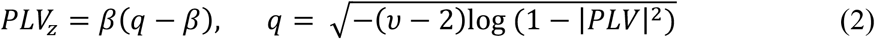

Here *z* indicates the rendered value, *β* is a constant 23/20, and *v* is the degree of freedom (product of trial and taper numbers).

### Identification of Synchronized Networks

The general approach of our network identification closely resembles the method described in (Hipp et al., 2011), and is only summarized below. An interaction between two cortical areas can be represented as a point in two-dimensional space and can be extended to four-dimensional space by adding the dimensions time and frequency.

Identifying networks of significant interaction is then equivalent to identifying continuous clusters within this high-dimensional space. In our approach, we first computed PLVs for all pairs of locations in all time and frequency bins and conditions with equation (1), which were later transformed with equation (2) to be Gaussian-like. Then t-statistics was performed across subjects, contrasting data after stimulus onset and baseline in all conditions. The resulting 4-dimensional matrix (location by location by frequency by time) was then thresholded with a *t*-value equivalent to *p* = 0.01, setting every connection to either 1 (connected) or 0 (not connected). Further, we performed a neighborhood filtering using a threshold of 0.5 to remove spurious connections: if the existing neighboring connections of a certain connection were fewer than 50% of all its possible neighboring connections, this connection would be removed. As one connection was defined by four numbers, representing its frequency, time, and the two locations (of a pair), two connections were defined as neighbors if they were identical in three of the four numbers, and the other value would only differ by one step, i.e., 100 ms in time, 0.25 octaves in frequency, or immediate neighborhood in source space. Two source locations were defined as immediate neighbors if they were directly connected in the regular polyhedron during source space construction. Eventually, the remaining neighbor connections formed the candidate networks for statistics. We defined their sizes as the integral of the *t*-scores across the volume. To evaluate their significance, random permutation statistics was performed to account for multiple comparisons across the interaction space. To this end, the network-identification approach was repeated 10000 times with shuffled condition labels to create an empirical distribution of network sizes under the null-hypothesis of no difference between conditions (Nichols and Holmes, 2002). The null-distribution was constructed from the largest networks (two-tailed) of each resample, and only networks with sizes ranked top 5% in these null distributions were considered as significant (i.e. *p* = 0.05) in the stimulus vs. baseline comparison. The final networks corresponded to cortical regions with different synchronization status among comparisons, and were continuous across time, frequency, and pairwise space. Applying this method, we compensated for the relatively arbitrary thresholding procedure during the generation of the 4-D connectivity array: the networks with fewer statistically stronger connections and networks with more statistically weaker connections were both considered.

### Further analysis of phase coupling networks!

Since we were interested in the contribution of single trials to the phase consistency, we calculated single trial phase locking pseudo values (STPLP) in addition to PLV. We did so with jackknife methods, similar to the calculation of single trial coherence pseudo values (Womelsdorf et al., 2006), according to:

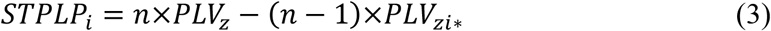

Here *i* is the single trial index, *n* the total trial number, and z indicates the rendered value (to make its distribution Gaussian); *i*\* indicates the trial group with the i-th trial left out.

Within the networks resulting from the contrast of stimulus and baseline, we computed STPLP for each trial. For the three conditions (T, V, and VT) separately, we then averaged the values across all location pairs, times and frequencies, as well as their corresponding values in the baseline interval. The data for all subjects were submitted to a 3 x 2 repeated-measures ANOVA, with two within-subject factors: stimulus condition (T, V, and VT) and brain state (stimulus vs. baseline). Both main effects and interactions were evaluated. Post-hoc analysis was performed to examine simple effects, when the interaction was significant.

As PLV might be vulnerable to volume conduction, we also computed the imaginary part of coherency as a complementary measure (Nolte et al., 2004). For all connections within our identified networks, we calculated the single trial imaginary part of coherency pseudo values (STICP), similar to the calculation of STPLP presented in equation (3). In order to confirm that our network identification was not driven by volume conduction, we compared STICP values using the same contrast as in our network identification approach.

Another possible confounding factor in connectivity analysis is the power of the oscillatory signals. As no coupling measure is independent of power, differences in power may cause differences in signal-to-noise ratio, which in turn may contribute to differences in coupling measures. Since we observed power differences between conditions in our data, we chose a subset of trials for each network by removing 5-15% of all trials (this ratio was fixed for all subjects in each network), to balance power across comparison groups. Then we redid the analysis on STPLP for all connections within the network and across the remaining trials to compare again between conditions of interest. Finding similar results as before would suggest that the effects were not driven by power differences.

### Phase-amplitude coupling

The above analysis was based on phase coupling between two locations in the same frequency. Another means of communication between brain areas is coupling across different frequencies: the next step of our analysis was to investigate cross-frequency coupling, or phase-amplitude coupling. When we had two central frequencies in a network in phase coupling, we further examined whether the phase of the slower frequency would modulate the amplitude of the higher frequency. For this purpose, the waveforms in the involved source locations of the phase coupling networks were first cut into bins of 250 ms length, centered around each time point, with a spacing of 100 ms (similar to analysis of phase coupling). Then in each bin, data were filtered for a certain frequency band according to the properties of the network, e.g., in the alpha (8 – 12 Hz) and beta (16 – 32 Hz) range. Next, the Hilbert transform was performed on these data, and the phase of the lower frequency signal (phase) as well as the amplitude of the higher frequency (amplitude) signal were extracted to compute the modulation index (MI, Tort et al., 2010). The phase information was binned into 18 intervals of 20°. The corresponding amplitude values for each bin were summed and then normalized by dividing by the mean across all bins. The MI was calculated as the Kullback-Leibler (KL) distance between the resulting distributions and the uniform distribution. The MI ranges from 0 (no modulation) to 1 (full modulation; please see also supplementary Figure S2 for a graphical illustration). We calculated MIs for each time point and between all source locations from network 1, whose connectivity values exceeded the average connectivity of all locations in the phase-phase coupling analysis. Then the differences between all time points and the baseline time point at – 200 ms were calculated and subjected to a network identification approach similar to the one for phase coupling, but with only 1 data point in the frequency dimension; thus, for the phase-amplitude coupling analysis three dimensions were considered: location of phase signal, location of amplitude signal, and time.

### Illustration of identified networks

To visualize networks identified by phase coupling, we projected them onto two subspaces separately. We integrated the *t*-values among connections over all spatial locations for each time and frequency bin to illustrate when and at which frequencies a network was active irrespective of the spatial distribution. Complementary to this, we also computed the integral of t-values in the connection space over time, frequency, and target locations for each location in the corresponding network. The result was then displayed on the template brain surface to reveal the spatial extent of the network independent of its intrinsic coupling structure and time-frequency characteristics. For illustration purposes, the data were interpolated from 300 to ~300,000 vertices on the inflated brain surface. The visualization of the phase-amplitude coupling networks was done similar to the phase coupling in the space extension. However, since the frequencies were predefined, we only plotted the one-dimensional interaction time course in the description of the dynamics. The spatial extent of the network was visualized in two separate projections on the brain surface, one for the phase signal locations and the other for the amplitude signal locations (see Figure 6A). The labeling of brain areas involved in our networks was based on their relative position to the gyri and sulci of the template brain (Fischl et al., 2004).

### Coupling and behavior

We also assessed the relevance of the connectivity networks observed with our clustering approach for participants’ performance on the behavioral task. For the phase coupling networks, we calculated the mean phase as well as power over all accurately answered trials of the data used for the network identification, assuming those values to be optimal for performance. Then the phase or power differences between every single trial (including also the wrong ones) and this optimal state were measured and the trials were sorted according to the median of the difference into two groups with identical trial number. This procedure resulted in a group with smaller difference to the optimal state and a second group with larger difference (for power and phase data). To examine whether phase or power information was related to participants’ behavior, we compared performance between the two groups. For the phase-amplitude coupling networks, we did not examine phase relations on a single trial basis. Instead, we correlated the strength of cross frequency coupling with behavioral performance across participants. Similarly, we also examined correlations between performance and power of both, the low and the high frequency signals of the coupled sites.

## Results

### Behavioral performance

Repeated-measures ANOVAs comparing accuracy and reaction time data for the different target cases revealed significant differences related to crossmodal stimulus congruence (F_2,28_ = 23.28, *p* < 0.0001 for accuracies; and F_2,28_ = 6.59, *p* < 0.01 for reaction times) with congruent VT targets being associated with the best performance (96.2 ± 3.6% correctly identified targets, mean ± standard deviation; reaction time was 376 ± 193 ms), followed by V (88.5 ± 9.2%; 390 ± 194 ms) and T targets (71 ± 16.3%; 411 ± 193 ms). Detailed analyses of the behavioral data have been reported elsewhere (Göschl et al., 2015).

### Phase coupling networks involved in target detection

After source reconstruction and spectral decomposition, we applied the data-driven network identification approach to the difference in PLVs between stimulus processing and baseline periods. Three highly structured networks associated with target detection were identified.

Network 1 (permutation-test, *p* < 0.0001, Figure 2A, B) was coupled mainly in the alpha (centered at ~10 Hz) and beta (centered at ~ 20 Hz) frequency range (Figure 2B). Synchronization decreased after stimulus onset compared to baseline, reaching a minimum at ~ 700 ms. Brain areas involved in this network were mainly located in the right hemisphere, contralateral to the response hand and the stimulated visual hemifield. The two most salient hubs within the network were supramarginal gyrus (SMG) and sensorimotor areas (Figure 2A). We use the general term ‘sensorimotor areas’ here, as the cluster included pre- and postcentral regions, as well as premotor areas, which were difficult to be discriminated from each other in our case. Next, we examined whether the connections within network 1 were mostly within a region or cross-regional. If the connections were random among the nodes, i.e., each node would have equal chance in connecting to any other node, either within its own region or to a node in another region, then we would expect about 47.7% of all connections to be cross-regional. Instead, in our data this ratio was 67.2%, indicating that the connections were more likely to be cross-regional than expected in a randomly connected network.

**Figure 2.**
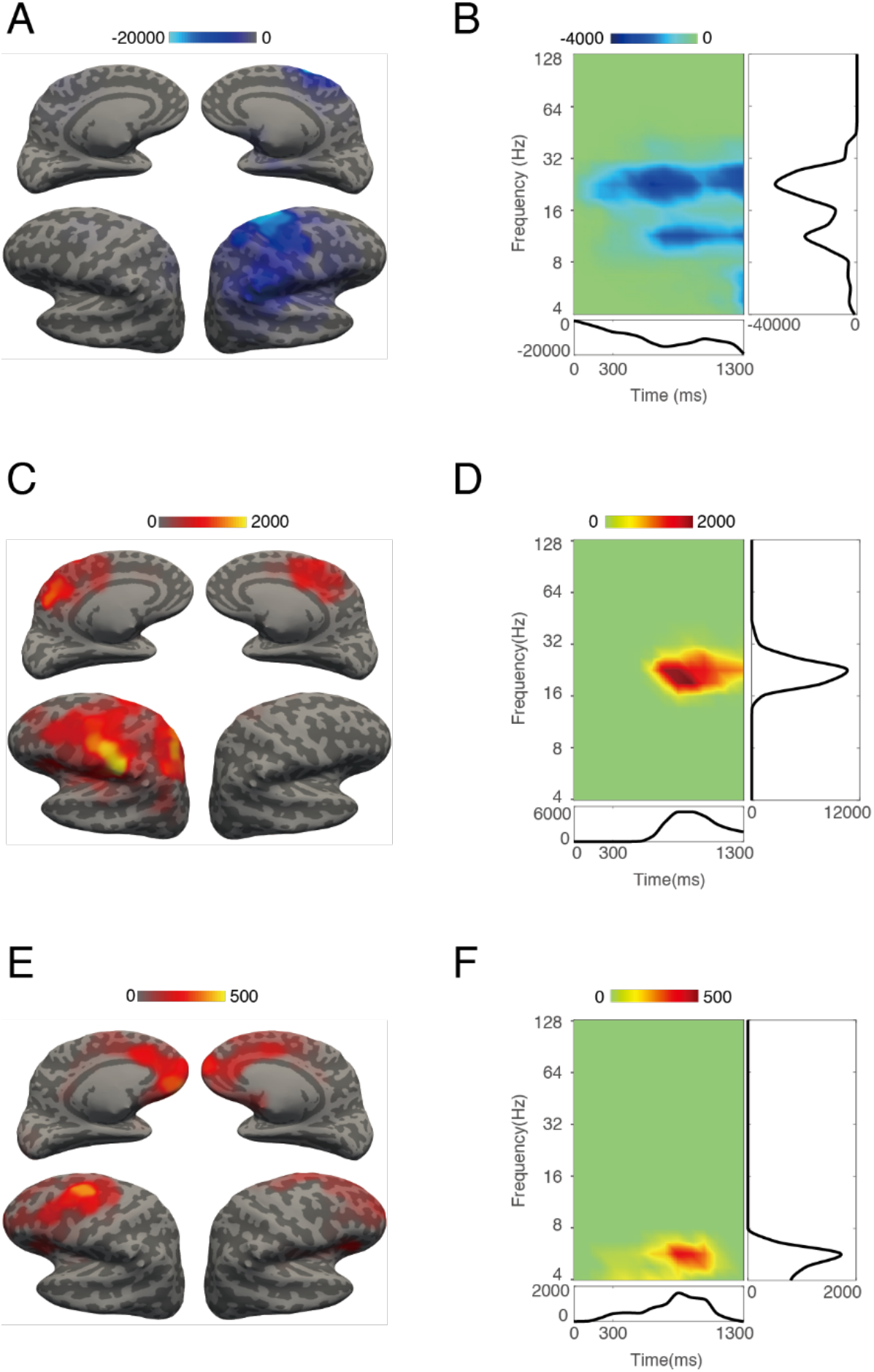
Phase coupling networks related to target detection. (A) Spatial localization of cortical areas involved in network 1. At each location, the color indicates accumulated t-values, which represent significance of PLV changes from baseline, between this location and all other locations, in all time and frequency bins. (B) Spectro-temporal profile of network 1, showing accumulated t-values over all location pairs showing significant changes in PLV from baseline at a given time and frequency point (color-coded). The line plots in the boxes right and below the time-frequency plot show the frequency and time accumulation of *t* values. (C - F) are similar to (A, B), but for network 2 and network 3.

Network 2 (permutation-test, p = 0.0016, Figure 2C, D) exhibited stronger beta band (centered at ~ 20 Hz) synchronization compared to baseline, starting at ~ 600 ms, and peaking between 800 – 900 ms after stimulus onset (Figure 2B). This network involved precuneus /midcingulate cortex in both hemispheres. The remaining activations were distributed in the left hemisphere only (Figure 2A), including superior parietal lobe, intraparietal sulcus, secondary somatosensory cortex, parietal ventral area, SMG, and precentral sulcus. The activation found in precentral sulcus may include more superior parts located in the frontal eye field as well as more ventral parts in the inferior frontal junction. Unfortunately, those regions were hard to separate spatially in our data, probably due to the limited number of sensors and the lack of individual anatomical MR images. Note that the tactile stimulation was delivered to the right index finger, matching with the somatosensory sources found to be part of this network. Again, we examined the proportion of connections within and across regions in the network. If the nodes connected randomly, we would expect about 79.5% of the connections to be cross-regional. In our data, we found 90.6%. of the connections to be remote instead. Thus, remote connections were predominant in network 2 as well, more than to be expected in a random network.

Network 3 (permutation-test, p = 0.0192, Figure 2E, F) was observed in the theta (centered at ~ 5 Hz) frequency range. The activation extended from ~ 200 ms to ~ 1200 ms after stimulus onset, peaking at ~ 800 ms (Figure 2F). Network 3 distributed mainly across frontal areas, including the upper part of the left PCS and the caudal middle frontal gyrus (which is potentially consistent with left frontal eye field). In addition, the anterior cingulate cortex and the orbitofrontal cortex in both hemispheres were involved (Figure 2E). The fraction of cross-regional connections within this network amounted to 85.7%, again higher than expected assuming random connectivity (63.0%).

Connectivity estimates such as coherence or PLV are vulnerable to artifacts from volume conduction: due to signal spread, sensors at two different locations can pick up a signal from the same generator, instead of representing two biologically coupled cortical regions (Nunez et al., 1997). The imaginary part of coherency, in contrast, excludes zero-phase lag coupling and thus is robust against effects of volume conduction (Nolte et al., 2004). To check for spurious coupling effects due to volume conduction in our analysis, we additionally calculated the imaginary part of coherency for all the connections in the detected networks. We then compared single trial pseudo values (STICP; see Methods for details) between stimulus and baseline and found significant effects, very similar to the ones reported for phase locking (see Figure 3A; p values are smaller than 0.01 in all three networks). This renders it unlikely that our results obtained using PLV were inflicted by volume conduction.

**Figure 3.**
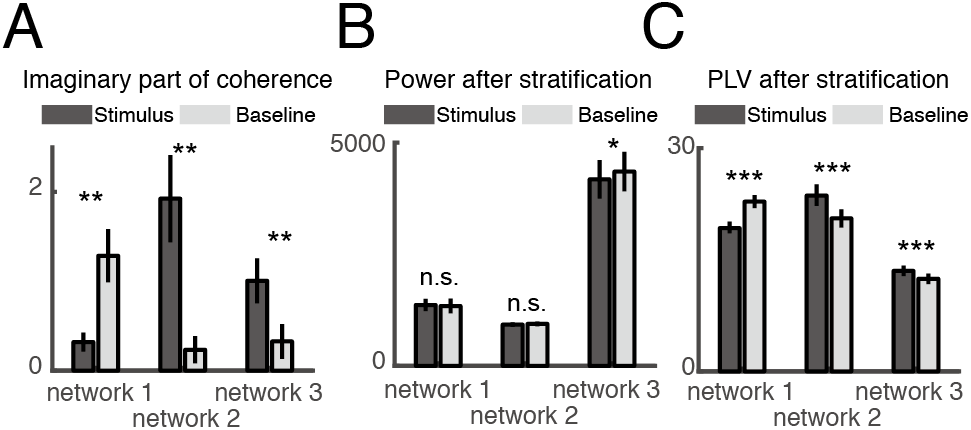
Further analyses of our three networks based on single-trial connectivity and power measures. (A) Comparison of single trial imaginary coherence pseudo values (STICP) between stimulus and baseline in all connections involved in the three detected networks. Error bars represent standard errors. (B) The comparison of power in the power-stratified trials. Note that in network 3, although the difference was significant, it was in the reverse direction as the difference in phase locking. (C) Comparison of single trial phase locking pseudo values (STPLP) between stimulus and baseline in all connections involved in the three detected networks, for power-stratified trials. * – p < 0.05; ** – p < 0.01; *** – p <0.001.

Another possible concern for the interpretation of coupling measures is power confounds. When two conditions differ in power, the connectivity in the condition exhibiting stronger power could be overestimated because of better signal-to-noise ratio. In addition to the analysis of long-range coupling, we therefore also tested local power effects in the involved brain areas at the corresponding time and frequency bins. We found significant differences in power, going in the same direction as the differences found in PLV. To rule out the possibility that the coupling effects were trivial consequences of power differences (caused by different signal-to-noise ratios), we applied the following steps: from the original data, we removed a small portion (5% to 15%) of the trials, such that the power difference between stimulus and baseline would either vanish (network 1 and 2), or even reverse (network 3; see Figure 3B). Next, the PLVs were compared again, yielding quite similar results to the original analysis (all p values of the comparison were smaller than 0.001; see Figure 3C).

### Phase coupling predicts performance

As shown in the above analysis (Figure 3B, C), we could be confident that the observed coupling effects were not mere consequences of differences in signal power. However, since these effects existed in parallel, the next question was which of the two measures, power or coupling, was more meaningful for brain function in our multisensory task? To evaluate the ecological relevance of our clustering results, we examined the relation of connectivity as well as power measures with performance: if one measure of neural activity could predict behavior better than the other, it should be more critical for the task at hand.

To evaluate the connectivity findings, we determined the phase delay between a pair of locations for every trial in the experiment. We defined the average of phase delays for each connection separately across all correct trials as the optimal delay (Figure 4A, red line in upper panel). Then, single-trial estimates were classified with respect to the optimal phase, either displaying a smaller difference (Figure 4A, magenta line in upper panel) or a larger difference to the respective optimal phase (Figure 4A, green line in upper panel). The networks detected in our analysis contained multiple connections, but by averaging the differences to the optimal delay over connections, we could evaluate the phase difference in a certain trial, as compared to its optimal status. All the trials (including correct and incorrect ones) were separated into two groups of equal size, one with smaller difference to the optimal phase, and the other with larger difference. Then, performance was compared between the groups for all three networks. As can be seen in Figure 4B, those trials with smaller difference to the optimal phase yielded better performance in network 2 and network 3 (Figure 4B). For network 1, a similar effect was observed, but only for the baseline period. We also grouped the data according to power instead of phase difference and compared performance, but no significant results were observed (Figure 4C). These findings suggest that coupling between remote brain areas (as mirrored in our phase locking clusters) was more relevant for task performance than local measures of signal power.

**Figure 4.**
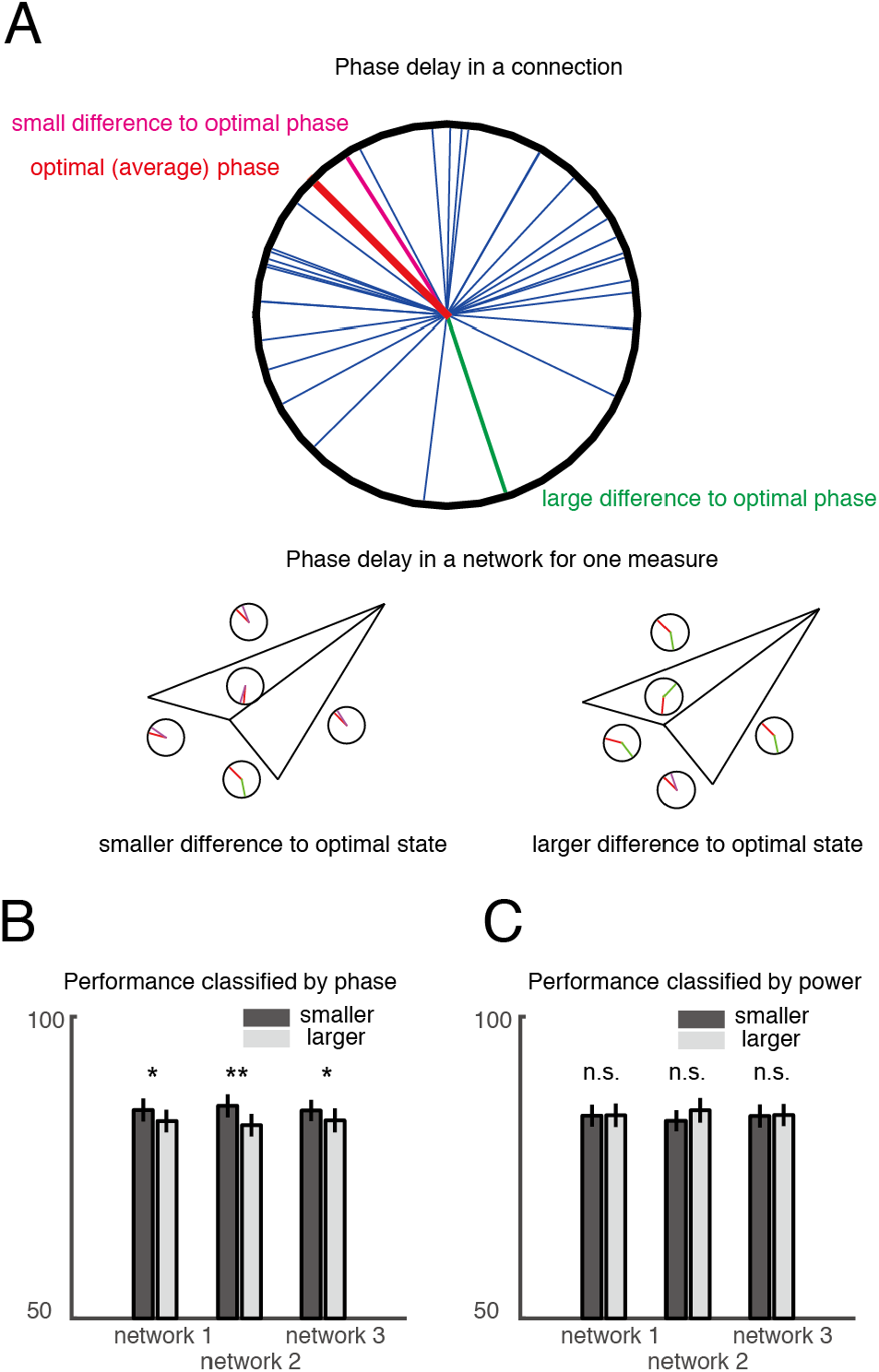
Phase delay predicts performance. (A) Top: Depiction of phase angles between a pair of locations in multiple trials. Each thin line in the circle represents the phase delay between two locations in a certain trial. The thick red line represents the optimal phase, defined as the average among all trials with correct responses. The magenta line represents the phase in a trial with smaller delay to the optimal phase; the green line represents a trial with larger delay to the optimal phase. Lower left: A trial where the connections to constitute a certain network have smaller delays to the optimal phase on average. Lower right: A trial where the connections to constitute the identified network have larger delays to the optimal phase on average. (B) Performance comparison between two groups of trials sorted by phase difference to the optimal phase. Note that the data shown for network 1 are from the baseline period whereas data for network 2 and 3 are from the stimulus period. (C) Performance comparison between two groups of trials sorted by power difference to the optimal power (defined as the average across all correct trials). * – p < 0.05, ** – p < 0.01.

### Phase coupling in network 2 is dependent on stimulus configuration

Follow-up analyses were conducted to test whether synchronization within the networks differed between stimulus conditions. We performed a repeated-measures ANOVA on accumulated STPLP in the network with two factors: stimulus condition (T, V and VT) and brain state (stimulus vs. baseline). In network 2, a significant main effect for the factor brain state (F_1,14_ = 35.08, *p* < 0.001, with Greenhouse-Geisser correction) as well as a significant interaction (F_2,28_ = 12.49, *p* < 0.001) were observed. The main effect for stimulus vs. baseline was expected since the network itself was identified based on this contrast. Next, we performed post-hoc analyses to examine differences in PLV between stimulus conditions. We found significant differences between the three stimulus conditions (F_2,28_ = 19.53, *p* < 0.001) during stimulus processing but not in the baseline interval (F_2,28_ = 1.30, *p* = 0.287). This indicated that coupling changed in response to different crossmodal stimulus configurations in network 2. Pairwise examination showed that the overall within-network phase locking was stronger in the congruent VT condition than the T condition (*p* = 0.002; see also Figure 5); and stronger in the V condition than the T condition as well (*p* < 0.001). Also, a subtle difference between the VT and V condition was apparent (*p* = 0.038). We did not observe any differences between stimulus conditions in network 1 and network 3.

**Figure 5.**
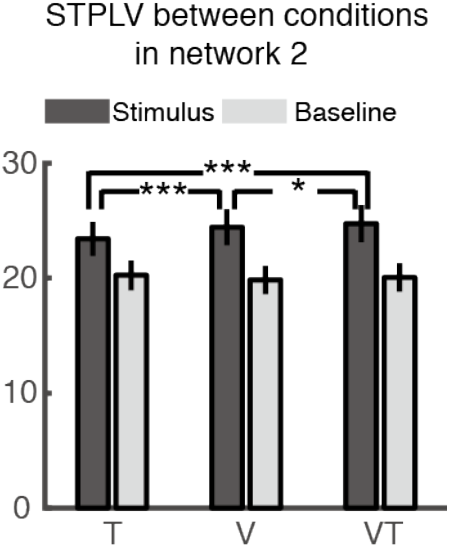
Comparison of single trial phase locking pseudo values (STPLP) across conditions in network 2. * – p < 0.05, ** – p < 0.01, *** – p < 0.001.

### Phase-amplitude coupling

Network 1 incorporated both alpha and beta frequency components, raising the question of whether and how the two might be related. One method to investigate interactions across different frequencies is to quantify phase-amplitude coupling (Tort et al., 2010). Here, we evaluated whether the phase of the slower alpha oscillation would modulate the amplitude of the beta oscillation. Based on the spectro-temporal signature of network 1, we chose frequencies between 8 and 12 Hz (alpha) and 16 and 32 Hz (beta) and a time interval from 600 to 1300 ms after stimulus onset for this analysis. Locations from network 1 displaying absolute *t*-values larger than average were chosen. A modulation index (MI, see Methods) reflecting the strength of phase-amplitude coupling was computed across all selected locations at the time bins described above plus the baseline bin 200 ms before stimulus onset. In addition, shift predictors were computed for these bins. To identify interactions across frequencies, we submitted MI values for the time interval of interest that both differed significantly from baseline and exceeded the stimulus vs. baseline difference in shift predictors to our clustering approach. One network (labeled network 4) was found. Network 4 was similar in spatial distribution to network 1, consisting of two clusters in SMG and sensorimotor areas (Figure 6A). Within this network, cross-frequency coupling was found to decrease relative to baseline. A negative correlation of MI values and performance (percentage of correct responses) was observed across participants (Figure 6C), i.e., participants who would more strongly reduce their alpha-beta phase-amplitude coupling in response to stimulation were more likely to perform well on the task. As a control analysis, we also examined the relation of both alpha and beta power at the locations involved in network 4 to task accuracy. Neither of the power measures showed a significant correlation with performance (Figure 6D, E). Also, we examined the relation of alpha– beta phase-amplitude coupling and response speed (reaction times). However, the significance of the correlation heavily depended on two participants, which is why we do not include this result here.

**Figure 6.**
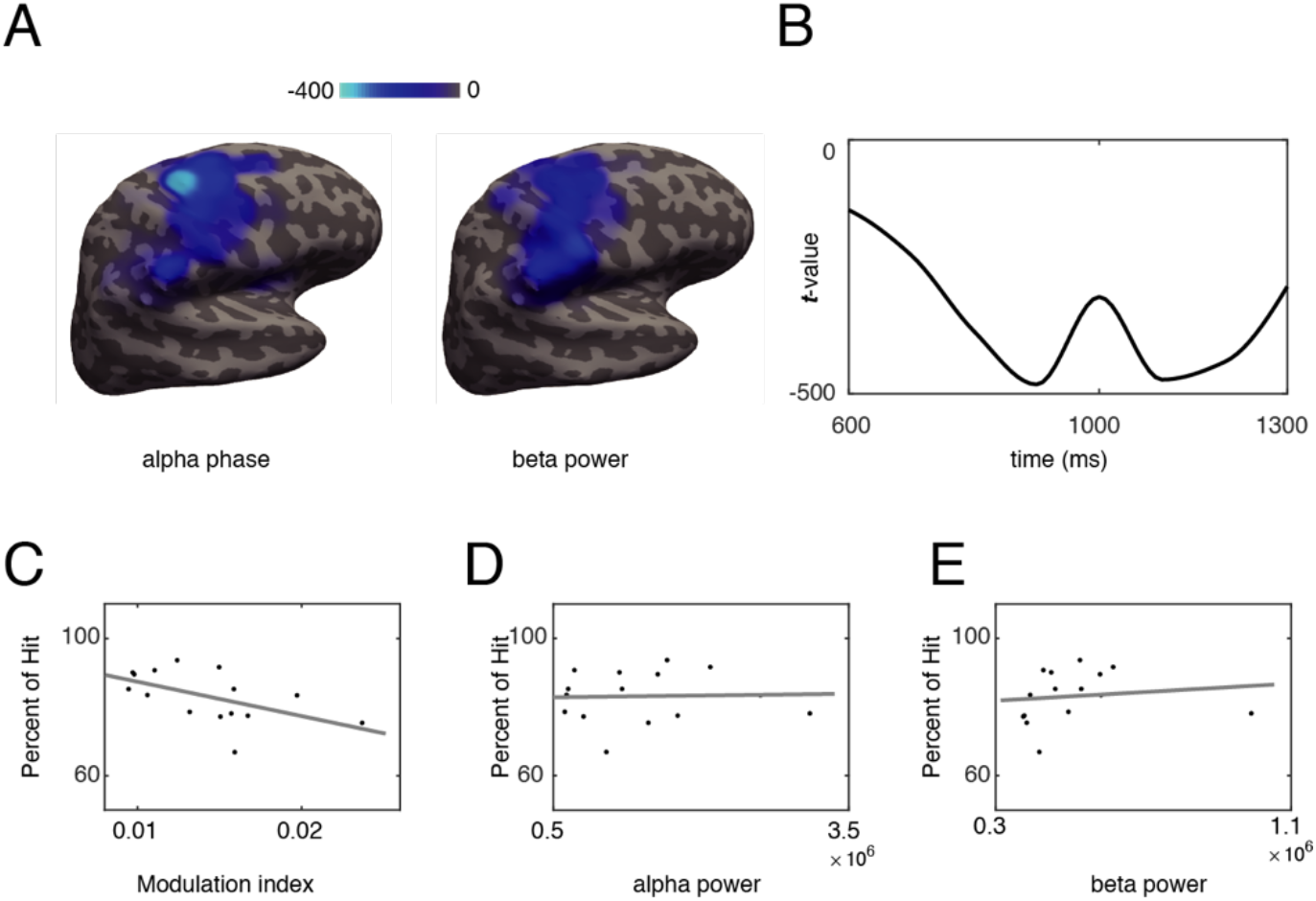
Alpha-beta phase-amplitude coupling relates to performance. (A) Spatial distribution of network 4. On the left, locations from which alpha phase modulates beta power are shown; on the right, locations where beta power was modulated are displayed. Accumulated t-values for the difference in phase-amplitude coupling between stimulus and baseline periods are shown color-coded. (B) Time course of the phase-amplitude coupling in network 4. (C) Correlation between accumulated modulation indices (MI) and task accuracy across participants. (D, E) Correlation of accumulated alpha/beta power and accuracy across participants. * – p < 0.05

## Discussion

In the present study, we re-analyzed a previously acquired EEG dataset (Göschl et al., 2015) with the aim to map functional networks mediating multisensory processes with a set of purely data-driven methods. The methodology employed in the current study allowed exploring neuronal dynamics in source space covering most of the cortex with a wide range of time and frequency being examined. This provided an evaluation of dynamic functional connectivity in cortical networks during a cognitive task without a priori constraints. Several networks of interacting sources were identified, which showed task- and condition-specific coupling effects across different frequency ranges, including the theta, alpha, and beta frequency band: (i) network 1 was characterized by phase-coupling in the alpha and beta band and comprised sensorimotor areas and supramarginal gyrus; (ii) network 2 showed phase coupling in the beta band and involved parietal and sensorimotor areas as well as precuneus/midcingulate cortex; (iii) network 3 was defined by phase coupling in the theta frequency range, comprising central, premotor, anterior cingulate and orbitofrontal areas; (iv) network 4 was characterized by alpha-beta phase-amplitude coupling and was similar in topography to network 1.

Our results strongly support the claim that neural synchronization is vital to long-range coordination in the brain, facilitating perceptual organization of sensory information and sensorimotor integration (Engel et al., 2001, 2013; Fries, 2005; Lachaux et al., 1999; Senkowski et al., 2008; Varela et al., 2001). This adds to growing evidence showing that large-scale cortical synchronization plays an important role in various cognitive functions, including selective attention (Buschman and Miller, 2007; Siegel et al., 2008), learning (Antzoulatos and Miller, 2014), and crossmodal integration (Maier et al., 2008). Especially connectivity in the beta-band was prominently represented in our findings, confirming central importance of this frequency range for the interaction between remote brain areas (Womelsdorf and Fries, 2007). The importance of the beta-band for long-range connectivity is also consistent with predictions from computational modeling (Kopell et al., 2000; Lee et al., 2013), evidence from invasive recordings in monkeys (Buschman and Miller, 2007; Pesaran et al., 2008; Saalmann et al., 2007), and clinical studies in different patient groups (Bangel et al., 2014; Leung et al., 2014; Uhlhaas et al., 2006).

The spatio-temporal profiles as well as the frequency characteristics of the networks identified with our data-driven analysis bare striking similarities to brain networks that have been studied extensively with other approaches: sensorimotor networks on the one hand, and executive control/attentional networks on the other hand. In the following we discuss the common grounds of these networks and the clusters found in our analysis, trying to embed our results within existing frameworks of sensorimotor decision making and attentional control.

### Suppression of long-range coupling in sensorimotor-SMG networks

Resulting from the contrast of stimulus processing versus baseline activity, network 1 exhibited suppression of phase coupling between remote brain locations in the right cerebral hemisphere, including sensorimotor areas (premotor areas as well as pre- and post-central structures) and SMG, located contralaterally to the response hand and the visual stimulation site. Network 1 as well as network 4 showed a spatial distribution and a frequency signature (centered around 10 Hz and 20 Hz) reminiscent of the well-studied mu rhythm (Chatrian et al., 1959; Pineda, 2005). The mu rhythm has been characterized by three properties (Crone et al., 1998; Thorpe et al., 2016): (i) topographic specificity: it mainly emerges in sensorimotor cortex; (ii) spectral specificity: it has an alpha (8 – 12 Hz) and a beta (15 – 25 Hz) component; and (iii) functional dependence on behavior: it decreases with motion-related activity. All three criteria are well met by our networks 1 and 4.

Besides its role in motor planning and execution, the mu rhythm has been studied in many other contexts and, amongst others, its role has been explored in brain-computer interface (BCI) applications (Wolpaw et al., 2002), social behavior and imagined movement (Carr et al., 2003; Cochin et al., 1999; Marshall and Meltzoff, 2011; McFarland et al., 2000; Oberman et al., 2005; Rizzolatti et al., 2001) and Parkinson’s disease (Devos and Defebvre, 2006). In the rich literature about the mu rhythm, it was usually the signal’s amplitude or oscillatory power that was mainly referred to. In our data, alpha- and beta-band power in sensorimotor cortex was reduced after stimulus onset and in preparation of the motor response, largely in line with earlier studies. Extending previous work, our study is among the first to describe the mu rhythm in terms of its coupling properties within and across frequencies, without pre-selecting time windows, cortical locations or frequency ranges. More importantly, our connectivity findings provided richer information as compared to spectral power. First, neuronal coupling directly related to participants’ performance. In network 1, we found trials with a smaller difference to the optimal phase to be more likely to be correct (within-subject effect). In addition, participants showing stronger ability to reduce alpha-beta phase-amplitude coupling during stimulus processing (defining network 4) performed better on the task. No such relation to behavior has so far observed with power-related measures. Second, our results not only showed coupling within sensorimotor cortex but also to other cortical sites, most prominently SMG. Studying these cross-regional connections of sensorimotor cortex to other brain regions might be a fruitful extension of the traditional understanding of the mu rhythm.

SMG – with a slightly broader spatial distribution often also labeled temporoparietal junction or TPJ (Schurz et al., 2017) – has been ascribed a number of different functions, one of them being the role of a hub in the ventral attention network for visual orienting (Corbetta et al., 2008). In previous fMRI studies on visual attention, activation in right TPJ were observed to increase for distractors with relevant features outside of attentional focus (Serences et al., 2005) but decrease for non-target objects in focused locations (Shulman et al., 2007). Also, it has been shown that activation in TPJ decreases with heavier visual short term memory load, inducing deficits in the detection of novel stimuli (Todd et al., 2005). In a recent fMRI study, hemodynamic activity was found to be negatively correlated with mu power in several brain regions, including sensorimotor areas and TPJ (Yin et al., 2016).

Coupling decreases in the alpha band between right-hemispheric SMG and sensorimotor areas paralleled our findings in the beta band. Alpha oscillations have most often been studied in occipital cortex, and a prominent theory holds that they mediate the gating of information between brain areas by inhibiting task-irrelevant regions (Jensen et al., 2012; Jensen and Mazaheri, 2010). Although alpha band activity in sensorimotor areas has traditionally been thought to be distinct from occipital alpha, there is recent evidence suggesting that sensorimotor alpha could also have inhibitory effects on task-irrelevant cortical sites and boost performance (Jensen and Mazaheri, 2010; van Ede et al., 2011). In our data, higher levels of alpha band synchronization between sensorimotor areas and SMG in the baseline period led to better performance, too.

Based on the findings presented above, the function of our network 1 might be conceived as follows: The prominent coupling between SMG and sensorimotor regions might reflect communication between visual attention areas and the motor system. During the baseline period, when participants anticipate the upcoming stimulus, enhanced alpha and beta connectivity both across and within sensorimotor cortex and SMG might indicate an inhibitory process. This inhibition might prevent the participants from being distracted in anticipation of the stimulus event. The behavioral significance of this connectivity pattern is illustrated by the finding that trials showing smaller delay to the optimal phase are more likely to be correct. After the presentation of the visuotactile stimuli, this inhibitory process might be released (Jensen et al., 2012; Jensen and Mazaheri, 2010), allowing the participant to detect the patterns and deliver the motor response accordingly. Our data suggest that the more effective the release from inhibition is achieved by a participant (mirrored in a reduction of alpha-beta cross-frequency coupling), the better his or her performance on the task turns out to be.

### Enhancement of long-range coupling in executive attention networks

Besides the orienting network, Posner and Petersen (1990, 2012) proposed the attention system to also include an alerting as well as an executive component. The executive network was proposed to be shared by different subsystems, and has usually been observed in target detection tasks (Posner and Petersen, 1990). In the following decades, studies primarily using fMRI identified brain substrates of the executive attention subsystem: first, a fronto-parietal network, which was associated with adaptive control and, second, a cingulo-opercular network, which was thought to mediate set-maintenance (Dosenbach et al., 2008). Our networks 2 and 3, obtained using a purely data-driven approach for clustering EEG source level connectivity data, well agree with these two networks both in terms of their spatial distribution and their function.

The brain areas forming our network 2 largely overlap with the fronto-parietal network, which has been described to consist of intraparietal sulcus, inferior parietal lobule, mid-cingulate cortex, dorsolateral prefrontal cortex and precuneus (Dosenbach et al., 2006; Dosenbach et al., 2007). This fronto-parietal network is believed to initiate and adjust task control within single trials in real time (Dosenbach et al., 2008). Consequently, trial-to-trial variations in stimulus validity (and differences in task difficulty between conditions as observed in our study) should yield activity differences within the network. This is exactly what we find when analyzing phase locking within network 2: differences between the T, V and VT conditions were apparent. Originally inferred from resting state fMRI data, the fronto-parietal network also seems to be important for active cortical processing during a cognitive task. We further showed that the phase information contained in our network 2 is predictive of performance.

Besides brain structures previously assigned to the fronto-parietal executive network, somatosensory cortex was also part of our network 2 – suggesting an interaction of attention and the processing of tactile information. In our experiment, participants were instructed to examine spatial patterns presented concurrently in the visual and the tactile modality and decide whether either of them was identical to a given target pattern. As inferred from detection accuracies and reaction times, this task was easier for visual than for tactile information. We propose that this difference in task difficulty might require enhanced communication between the executive attention network and somatosensory cortex – and be the reason why we do not observe significant connectivity to visual areas. It seems as if our participants’ attentional resources were not equally distributed across the two task-relevant sensory domains but instead enhanced connectivity for visual target detection led to better performance. Considering that our experimental task demanded the processing of tactile *and* visual information, it remains an open question why connectivity in network 2 was mainly established between attention-related structures and somatosensory cortex. The target detection design realized in the current study does not allow a clear separation of behavioral and brain responses to tactile and visual input – complicating the interpretation of the connectivity findings, especially the differences in coupling between conditions. Follow-up work should remedy this shortcoming, e.g., by including task conditions of selective unimodal attention as well as unimodal (visual or tactile only) sensory stimulation.

As all the stimuli were presented at expected timing and known locations, we refrain from interpreting our network 2 as the intensively studied orienting network, although the involved brain areas largely overlapped (Corbetta et al., 2008). However, it would be promising to further investigate the relation of the orienting network and the executive network with our data-driven methods in future studies. In our case, network 2 was mainly located in the left hemisphere, prominently involving somatosensory cortex (in line with the fact that the right index finger was stimulated). However, previous reports (Ruff et al., 2009) showed visual attention networks (involving frontal eye fields and intraparietal sulcus) to be lateralized to the right hemisphere. To resolve this spatial ambiguity and isolate attention- and stimulus-related processing, it would be interesting to examine the distribution of the network depending on the site of tactile (and also visual) stimulation in future work.

Alongside the fronto-parietal network, the cingulo-opercular network – including the anterior cingulate cortex, anterior prefrontal cortex, anterior insula and frontal operculum – has been described to keep a stable set-maintenance over the entire task epoch (Dosenbach et al., 2006; Dosenbach et al., 2007). Indeed, our network 3 showed high spatial overlap with this cingulo-opercular network. The fact that we do not find differences between conditions within network 3 seems to support the putative main function of the cingulo-opercular network to mediate a stable set-maintenance, in our case to represent the target pattern. Theta oscillations have also been reported during working memory tasks in human neocortex (Raghavachari et al., 2001; Roux and Uhlhaas, 2014) and are commonly associated with maintaining relevant information. These observations are well in line with our interpretation of network 3 supporting a stable set-maintenance – a process probably making use of working memory capacities. The absence of sensory areas in network 3 might point to an abstract pattern maintenance or storage– independent of the sensory modality involved. Also, network 3 could be confirmed to be functionally relevant: closer relation to the optimal phase led to better performance.

During the processing of crossmodal information, our networks 2 and 3 showed enhanced long-range neuronal coupling between remote areas that have previously been associated with the fronto-parietal and the cingulo-opercular networks of attention. In addition, we demonstrated the relevance of these brain networks for behavior by relating phase information and detection performance. Though derived with a different methodology (EEG vs. fMRI) and during a multisensory detection task (rather than at rest), our networks 2 and 3 can be well interpreted within the framework proposed by Posner and Petersen (1990, 2012). In the current study, we did not manipulate attention experimentally. Nonetheless, our data-driven approach revealed clusters of functionally connected brain areas strongly resembling attention networks observed in fMRI data – making this method potentially interesting for future attempts to dissociate different attentional subsystems.

### Methodological considerations

The time-varying power of an oscillating signal marks first-order changes and allows only indirect inference regarding long-range interactions among distributed sources (Schneidman et al., 2006). In the current study, we sought to explicitly assess second-order changes, namely pairwise phase dynamics of different brain areas, quantified as either phase or phase-amplitude coupling. Only recently, the idea of single rigid brain modules specialized for certain functions has been extended in favor of adaptive and flexible brain networks favoring collaboration among multiple brain sites. Much of this research (e.g. Friston, 2002; Sporns et al., 2004) was done using fMRI – a technique benefiting from its high spatial resolution and its ability to provide a comprehensive picture of whole-brain connectivity patterns. However, the BOLD signal is an indirect measure of neuronal activity and sluggish in its temporal resolution (Ogawa et al., 1990). Invasive depth electrodes in contrast measure neuronal responses directly, but only allow to study a small fraction of brain tissue. EEG and MEG are characterized by both precise timing and a reasonable spatial coverage. We therefore believe that the analysis of EEG/MEG source level connectivity data using data-driven and constraint-free approaches like the one presented here helps to extend our understanding of the dynamic nature of brain networks at rest and during cognitive processing.

In our analysis, both phase coupling and phase-amplitude coupling were quantified. To rule out confounding effects of signal strength, we examined effects with stratified trial numbers and controlled for shift predictors in the phase-amplitude coupling analysis. The question how phase coupling and phase-amplitude coupling found in our data relate to each other is still open. At one cortical location, there may be various neuronal groups oscillating at different frequencies, e.g. high and low gamma activity have been shown to co-exist in different layers of the same cortical site (Ainsworth et al., 2011). Due to the limited spatial resolution of techniques like EEG and MEG, it is hard to distinguish activity from different cortical layers (though there are promising attempts to isolate signals from deep and superficial laminae using MEG; see for example Bonaiuto et al., 2018; Troebinger et al., 2014). Thus, in principle, it is possible that a local group of neurons oscillating in the alpha frequency range couples with a remote group within the same frequency, and then the target neurons locally modulate beta oscillations (via local phase-amplitude coupling); alternatively, it could also be that one neuronal group oscillating in alpha frequencies directly modulates (via cross-regional phase-amplitude coupling) a remote group oscillating in beta frequencies, while long-range coupling within same frequencies could take place in parallel. The interplay of neuronal coupling across and within frequencies and their functional relevance demands further investigation. However, our results of coupled cortical networks and their significance for behavioral performance further support the idea that oscillations are crucial for information transfer in the brain (Engel et al., 2001; Engel et al., 2013; Fries, 2015).

Unlike previous work (Hipp et al., 2011), we did not observe a dissociation between local oscillatory activity and long-range synchronization but in contrast found parallel effects in power and phase coupling measures. Our additional analysis minimized the risk that the observed coupling patterns were trivial technical consequences of the power differences. After stratifying the trials to remove or even inverse the power relation, the coupling differences were still well preserved. Further analyses suggested that coupling properties, but not power, could predict behavioral performance. However, it is likely that they are functionally related. One hypothesis is that successful communication between two remote brain areas also enhances local activities, i.e. the local firing patterns would be more synchronized – yielding stronger local power measured on the scalp. Given datasets of sufficient size, this hypothesis could be tested in future work using causality measures.

Of course, the method employed here has its limitations, too. One constraint refers to the spatial resolution of our clustering approach: in order to deal with the high-dimensional data space and to avoid extensive computational effort, we chose a relatively coarse resolution for the analysis in source space. However, the spatial resolution was compatible with our anatomical data: instead of individual magnetic resonance images (MRI) and individually measured sensor locations, we used template brain information and averaged sensor locations. Another limitation concerns the sensitivity of our method. Exploring data in a high-dimensional space including multiple time points, cortical locations and frequencies implicates accounting for a massive multiple-comparison problem – making the approach less sensitive. It is therefore possible that some (subtler) connectivity effects failed to be identified with our network identification approach, for example involving early visual areas or frequency components in the gamma range. !

## Conclusion

In the current study, we provide evidence for a functional role of long-range neuronal coupling in integrating distributed information in the human brain. Our data suggest that cortical networks defined by synchrony mediate crossmodal sensory processing, sensorimotor transformations, as well as executive control and set maintenance. Importantly, we demonstrate that inter-areal synchronization predicts behavioral performance, illustrating the functional relevance of large-scale coupling for cognitive processing.

## Acknowledgments

This research was supported by grants from the German Research Foundation (SFB 936/A3/B6, TRR169/B1/B4) and the European Union (ERC-2010-AdG-269716) awarded to A.K.E. and P.K. The authors thank Guido Nolte, Jörg Hipp and Till Schneider for methodological support, Jonathan Daume for constructive discussions and Julia Diestel for assistance in data recording.

---

**Figure S1.**
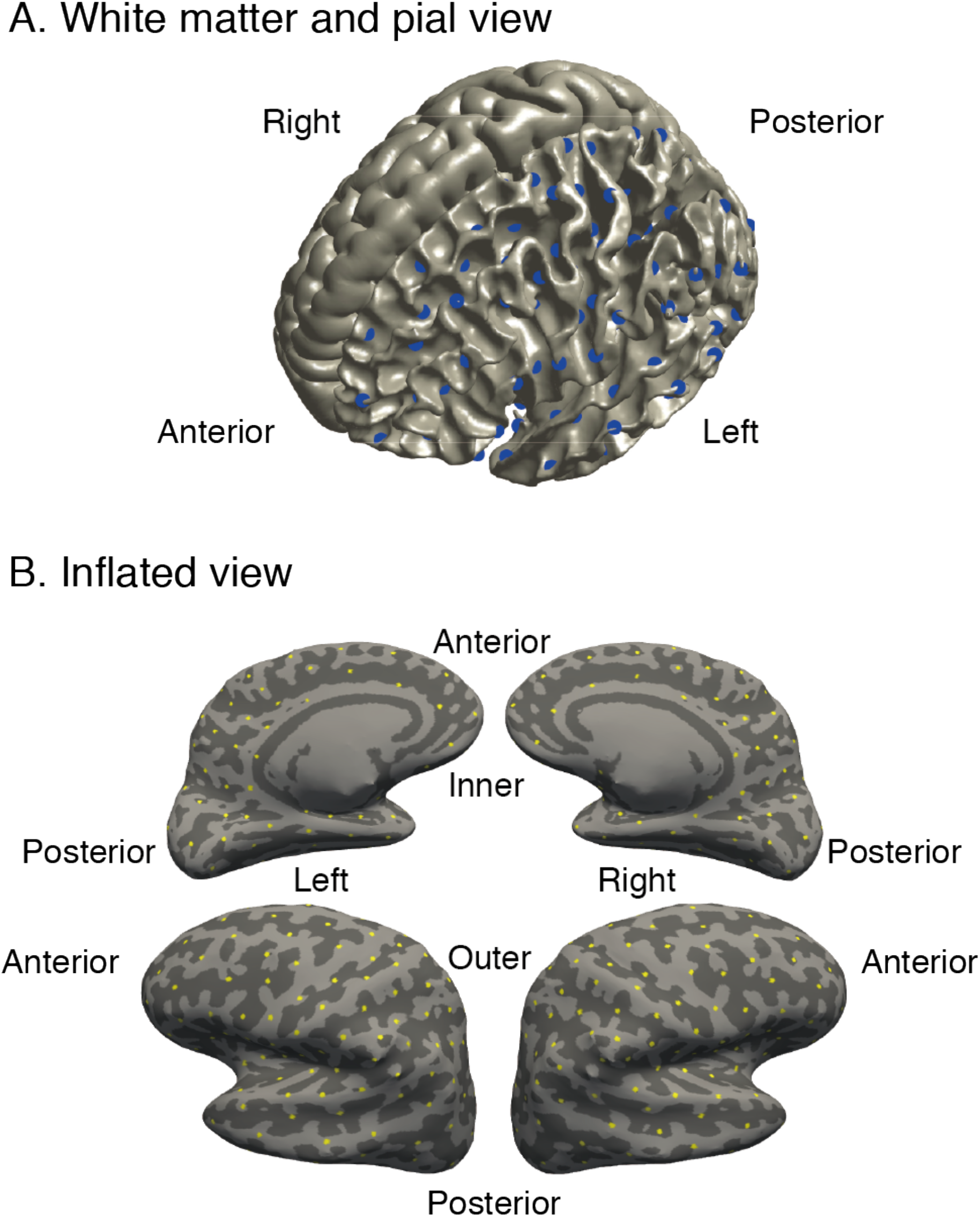
Definition of the source space. (A) The white matter (left hemisphere) and pial (right hemisphere) view of the brain template. The source locations were evenly distributed on the surface of the white matter, as indicated by the blue dots in the left hemisphere. (B) The inflated view of the brain template. The source locations are indicated by the yellow dots on the inflated surface.

**Figure S2.**
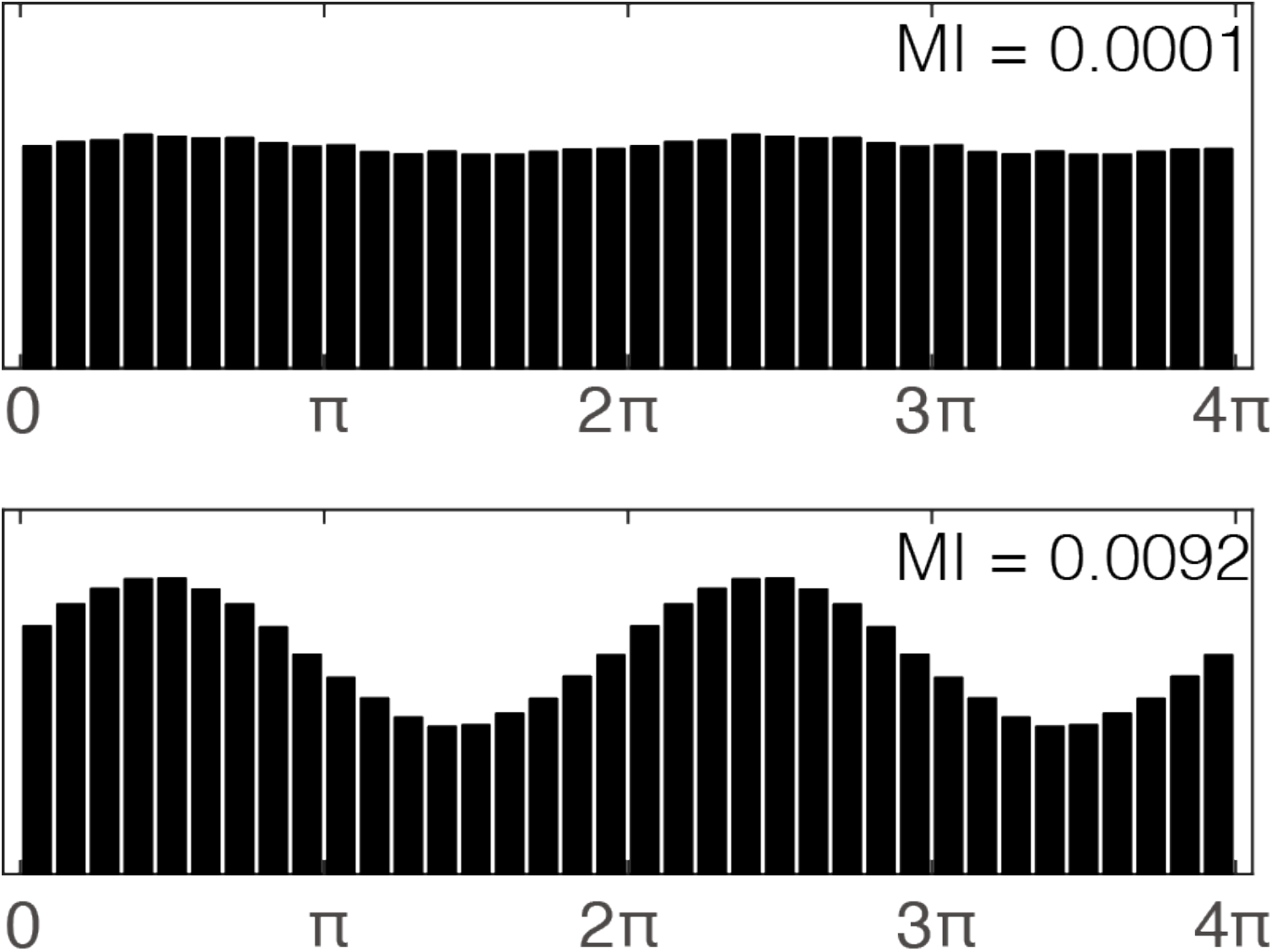
Illustration of phase-amplitude coupling displayed using the modulation index (MI). The bars on the y-axis indicate the amplitude of high frequency components, as modulated by the low frequency phase (shown on the x-axis). The plot in the lower panel exhibits stronger phase amplitude coupling, as indicated by higher MI.

